# Development and Validation of an LC-MS/MS Based Quantitative Assay for Marmoset Insulin

**DOI:** 10.1101/2021.06.24.449777

**Authors:** Robinson W. Goy, Hemanta K. Shrestha, Toni E. Ziegler, Natalie J. Dukes, Ricki J. Colman, Amita Kapoor

## Abstract

**Introduction:** Insulin is a peptide hormone that is secreted by the β cells of the pancreas and is essential to the metabolism of carbohydrates, fats, and proteins in the body. The marmoset insulin peptide is not homologous with human insulin and therefore commonly available assays do not work for this species. Due to the increasing popularity of marmoset research, a simple, specific assay for the quantitation of marmoset insulin is needed. This study aimed to develop and validate a bottom-up proteomic workflow with trypsin digestion and analysis using LC coupled with triple quadrupole mass spectrometry (LC-MS/MS).

**Methods:** Marmoset serum proteins were subjected to denaturation, reduction, and enzymatic cleavage to extract a unique, 7 amino acid peptide for quantitation of marmoset insulin.Resolution of the peptide was achieved by LC-MS/MS using electrospray ionization operating in positive mode. Calibration was achieved by aliquot dilution of fully synthetic marmoset insulin tryptic peptide into macaque serum. A stable-isotope labeled (^13^C, ^15^N) synthetic marmoset insulin tryptic peptide was used as internal standard.

**Results:** The assay was fully validated according to bioanalytical method validation guidelines for linearity, precision, and dilution linearity using purified marmoset insulin. The limit of detection was 4.28 pmol/L and the limit of quantitation was 8.62 pmol/L. Biological validation was achieved by comparison of samples previously run by radioimmunoassay and measurement of the marmoset insulin response to glucose via an oral glucose tolerance test (OGTT). The physiological range of marmoset insulin was shown to be 84.5 to 1222 pmol/L.

**Conclusions:** This paper presents a simple, reproducible method to measure marmoset insulin using LC-MS/MS.

## INTRODUCTION

The common marmoset (Callithrix jacchus) has been used in biomedical research for decades [1]. Within the last 10 years, popularity of this model has increased dramatically, prompted to a large degree by awareness of their utility for aging, neuroscience, metabolic disorders, and transgenic and genomic editing research [2], [3]. Many factors make the marmoset an attractive model system including their genetic and physiological similarity to humans, relatively short lifespan, high fertility, rapid development, small size, and human-like social structure [4]. However, the lack of marmoset-specific biomarker assays has hampered the use of marmosets as models.

Metabolic syndrome is a cluster of biological factors characterized by abdominal obesity, dyslipidemia, hypertension, and type 2 diabetes mellitus (T2DM) that are associated with increased risk of multiple chronic diseases [5], [6]. Marmosets are a good model for metabolic syndrome as they exhibit metabolic dysfunction associated with high body weight and aging by six to eight years of age [7], [8]. Insulin, a peptide that is secreted by the pancreatic β cells, is a biomarker for metabolic syndrome. Its main function is to maintain optimal blood glucose levels by facilitating glucose uptake by cells. Due to this role, high fasting insulin and an impaired insulin response to glucose levels are some of the hallmarks of metabolic dysfunction [9], [10].

The insulin peptide sequence is highly conserved across most species only differing in one to four positions with 92–98% identity [11]. While the marmoset pancreas exhibits the same structure, marker genes, and the presence of glucose transporters as human pancreas [12], the marmoset insulin peptide sequence differs in 5 amino acids from human and old world primates rendering limited or no immunologic cross-reactivity [13], [14].

Insulin is typically measured with immunological techniques, historically beginning with radioimmunoassay (RIA) and now with immunochemiluminometric assays on automated platforms [15]. Ziegler et al. [16] fully validated an immunoassay to measure marmoset insulin using a polyclonal porcine insulin RIA where the antibody cross-reacted 100% with human insulin, but was less specific than the human insulin RIA (Millipore Corp., Billerica, MA). However, in 2016, the antibody was depleted and the new one no longer cross-reacted with marmoset insulin. Our lab screened numerous commercially available immunoassays, including bead-based multiplex kits, to determine if an antibody used for insulin measurement in other species would cross-react with marmoset insulin, but this approach was unsuccessful. We considered creation of a custom immunoassay but it was logistically difficult. A relatively large amount (1-2 mg) of endogenous, pure analyte is required to create antibodies. Marmosets are in scarce supply due to their popularity as a research model, and being small monkeys with low blood volume, marmoset insulin is not commercially available, making the immunoassay approach not feasible.

We opted to develop a method to measure marmoset insulin using a bottom-up tryptic peptide LC-MS/MS approach. To our knowledge, there are currently no assays for any analytes in marmosets utilizing this technique. Since the marmoset insulin peptide sequence was published [13], we synthesized the tryptic peptide that was appropriate for quantitation [17]. Others have developed methods using LC-MS/MS for measurement of human insulin and other animal insulins. Their techniques include quantitation using B-chain [15], enrichment via immunocapture [18], and intact synthetic insulin analysis [19].

In this paper, we present the development and validation of a novel LC-MS/MS based method to quantify marmoset insulin utilizing a species-specific tryptic peptide. Utilization of this technology ensures independence from commercial sources.

## MATERIALS AND METHODS

### Animal care

Marmosets used for validation in this study were housed at the Wisconsin National Primate Research Center (WNPRC) at the University of Wisconsin-Madison (UW-Madison). The study reported here was conducted with institutional animal care and use committee (IACUC) approval and in accordance with national standards on animal welfare.

### Standards and reagents

HPLC grade water, ammonium hydroxide, and glacial acetic acid were purchased from Fisher Scientific (Waltham, MA); HPLC grade acetonitrile and methanol were purchased from Honeywell (Charlotte, NC); MS grade trypsin, iodoacetamide (IAA), high quality trifluoroacetic acid, and low protein binding microcentrifuge tubes were purchased from Thermo Fisher Scientific (Waltham, MA); Ammonium bicarbonate, dithiothreitol (DTT), and formic acid were purchased from Sigma-Aldrich (St. Louis, MO). Fully synthetic marmoset insulin tryptic peptide (MITP) was purchased from Genscript (>98% purity) (Piscataway, NJ). Fully synthetic, isotopically labeled insulin tryptic peptide (MITP-IS) was purchased from Thermo Fisher Scientific (>95% purity). All protein components were fully characterized by HPLC and MS by the manufacturers.

### Preparation of calibrators, controls, and internal standards

A stock solution of MITP was prepared at 6 µmol/L in water and stored at -80°C until use. Immediately before the assay, the stock solution was thawed and diluted to a working solution in 50 mM ammonium bicarbonate buffer, pH 8.0, to a final concentration of 3.45 nmol/L. The working solution was aliquot diluted into macaque serum to generate an 8-point calibration series (i.e., 3100, 1550, 776, 388, 193.4, 96.8, 48.4, and 24.2 pmol/L). Macaque serum was used as a blank matrix for this method since the insulin fragment is unique to marmosets and macaque serum does not contain native marmoset insulin. This serum came from quality control (QC) material that was used for other methods in the lab [e.g. 20]. QC pools for marmoset serum were collected and stored at -80°C in 150 µl aliquots. The isotopically labeled MITP used as internal standard (IS) was prepared in water at a concentration of 250 nmol/L. Single use aliquots were stored at -80°C and thawed before use. The IS was added to the calibrators, controls and serum samples after protein precipitation (See sample preparation below).

### Preparation and purification of intact marmoset insulin

Briefly, marmoset insulin was purified from pancreas using a modification of published methods [14]. Pancreas was acquired from the WNPRC pathology unit when marmosets were at necropsy (for other studies or vet-directed). It was thawed, weighed, cut into small pieces, transferred to a tube containing 5 volumes of cold acidic alcohol (0.2 M HCl in 75% EtOH) and ground with a Polytron grinder. After overnight incubation at -20°C, the mixture was centrifuged and the supernatant was collected. The supernatant was then precipitated, dissolved in 10 ml of 1 M acetic acid, dried, and then reconstituted with 100 µl of acidified water and stored at -80°C until purification. The marmoset insulin was purified by HPLC (Vanquish, Thermo Fisher) using reverse-phase C18(2) column (Luna 5 micron 250 mm x 4.6 mm, Phenomenex) and eluted with a linear gradient of 25–40% acetonitrile (0.085% trifluoroacetic acid). The fraction containing the diode array detected (214 nm) insulin peak was dried in Speedvac, reconstituted with 100 µl of acidified ddH2O, and stored at -80°C. Confirmation that the fraction contained only the purified insulin peak was conducted with MALDI-TOF at the University of Wisconsin-Madison Biotechnology Center, Mass Spectrometry Facility.

### Sample preparation

Serum from unknowns (used for biological validation, below) and QC pools were thawed, vortexed, and 50 µl of serum was transferred to a low protein binding 1.5 ml microcentrifuge tube. Large proteins were precipitated by a 1:1 addition of 1/49/50 acetic acid/methanol/acetonitrile solution. Samples were adjusted to pH > 9 by addition of a 5% ammonium hydroxide solution and allowed to incubate at -20°C for 30 min. The resulting precipitate was centrifuged for 10 min at 13,000 rpm. Macaque serum was precipitated the same way. The supernatant was transferred to a clean low protein binding microcentrifuge tube and 10 µl IS was added. Intact marmoset insulin and the MITP used for calibration was added to protein precipitated macaque serum at this step. To denature the native insulin, 50 µl of 0.05% RapiGest SF surfactant (Waters Corporation) was added and allowed to incubate at 80°C for 10 min. Disulfide bonds were broken by adding DTT to a final concentration of 20 mM and incubated at 60°C for 20 min. The resulting free cysteines were alkylated by addition of IAA to a final concentration of 40 mM followed by an incubation at room temperature, in the dark, for 30 min. The peptides were then enzymatically cleaved by addition of trypsin (Thermo scientific) and incubated at 37°C with mild agitation overnight. The trypsin reaction was stopped by raising the pH to >4 and incubated at 37°C for 30 min.

Sample cleanup was tested using Oasis MCX 1cc solid phase extraction (SPE) cartridges and Oasis MCX 96-well µelution SPE plate (Waters corporation). For the 1cc MCX SPE cartridges, MITP was eluted in 2 column volumes of 5% ammonium hydroxide in methanol. The resulting eluate was dried via vacuum centrifugation and reconstituted in 30 µl 20% acetonitrile in water with 0.1% formic acid. For the MCX µelution plate, MITP was eluted twice with 25 µl 5% ammonium hydroxide in methanol, and the resulting eluate was diluted 1:1 with HPLC grade water for analysis.

### Analytical chromatography and MS conditions

Samples were analyzed on a QTRAP 5500 triple-quadrupole linear ion trap mass spectrometer (Sciex) equipped with an electrospray ionization (ESI) source. The system included two Shimadzu LC20ADXR pumps and a Shimadzu SIL20ACXR autosampler. A sample of 20 µl was injected onto a XSelect CSH C18 XP column (2.1 mm x 100 mm, 2.5 µm, 130Å) (Waters) analytical separation was achieved using a mobile phase consisting of water with 0.1% formic acid (Solution A) and acetonitrile with 0.1% formic acid (Solution B), at a flow rate of 200 µl/min. MITP was eluted by a linear gradient of 15% to 25% solution B over 5 min. Solution B was increased to 100% over the next 0.1 min and held for 0.5 min, followed by a decrease back to starting conditions of 15% solution B over 0.1 min and held for 3 min for a total run time of 8 minutes.

### Method Validation

The developed method was validated for the calibration curve performance and precision in determination of insulin in marmoset serum according to bioanalytical method validation guidelines of the US Food and Drug Administration [21].

#### Linearity, limit of detection (LOD), and limit of quantitation (LOQ)

A calibration curve was developed by multiplying the ratio of the peak area of the calibrator to the peak area of IS and the concentration of the calibrator in pmol/L. Linearity was evaluated by monitoring coefficient of determination (r2) of the calibration curves. The performance of the assay was characterized by determining the LOD and the LOQ. QC pools were serially diluted and measured 5 times to calculate the concentration at which the CV was no higher than 20% and the LOD was calculated from the marmoset pool dilution which could be measured at a CV > 20%. The LOD and LOQ were also measured in 5 replicates in calibrators spiked in macaque serum. *Precision*. Precision was evaluated by measuring the coefficient of variation (CV) of the measurements of the high and low QC pool. QC data was collected at two levels (whole marmoset serum “high QC” and 1:1 diluted marmoset serum “low QC”). Values for the QC pools were collected in 20-25 determinations over the course of 2 months in separate batches.

#### Dilutional linearity

A portion of the purified insulin was diluted to demonstrate dilutional linearity and to confirm that the method developed was measuring intact marmoset insulin.

### Biological Validation

#### Oral glucose tolerance testing (OGTT)

OGTT was conducted in 6 adult marmosets to demonstrate their insulin response to dextrose. These marmosets were part of a larger study to understand the role of dietary fat on brain development so a wide physiological range of glucose and insulin were expected. Subjects consisted of two females and four males between the ages of nineteen and twenty-five months. The details of housing and husbandry are published [22]. Dextrose volume was based on marmoset weight (50% dextrose solution at 5 mL/kg). A 0.5 ml baseline blood sample was collected to measure starting glucose and insulin concentration after which, monkeys were administered the calculated volume of dextrose orally. Blood draws were then taken at 15, 30, 60, and 120 minutes. Glucose was measured with a glucometer (YSI 2900D). Blood samples were processed for serum and stored at -80°C.

#### Method comparison

Twenty-four marmoset serum samples (stored at -20°C) that had previously been run for measurement of insulin and had sufficient volume (50 µL) remaining were tested with the newly developed method. These were the last samples that Assay Services ran before the manufacturer changed the antibody of the validated porcine insulin RIA and they were stored with the intention of being used to validate a new marmoset insulin assay.

### Data analysis

Analyst software version 1.6.2 was used for all LC-MS/MS data acquisition and analysis. All statistical analyses were performed using Analyst and Prism version 8.4.3. For the OGTT, glucose and insulin levels were analyzed by one-way repeated measures analysis of variance (ANOVA) (time) with Tukey’s multiple comparison test and was qualitatively analyzed by determining the percent change from baseline for glucose and insulin. The comparison between samples run by RIA and the new LC-MS/MS method were analyzed using Pearson’s correlation analysis. The calibration curve and the intact marmoset dilution series were analyzed with linear regression.

## RESULTS

### Optimization of sample preparation

Trypsin concentration was optimized by addition of 0.1, 0.2, 0.3, 0.4 and 0.5 µg in triplicate. Area counts of marmoset insulin plateaued with 0.3 µg, therefore that concentration was chosen for all further experiments. The digestion time was 16 hours based on manufacturer recommendations and consideration of lab logistics.

Strong cation exchange mixed-mode (MCX) SPE was chosen for sample cleanup and desalting and tested in cartridge and µelution plates form. The cartridge required sample dry-down before reconstitution and injection which led to protein adsorption and inconsistent results. Protein precipitation also occurred resulting in poor chromatography and increasing backpressure. For these reasons, the 96-well µelution MCX SPE plate (Waters corporation) was chosen for sample clean up as the dry-down step was not required.

### Optimization of the tryptic peptide, mass spectrometry, and chromatography conditions

The marmoset insulin tryptic peptide (GFFYAPK) was selected for quantitation based on uniqueness, ease of synthesis, and multiple reaction monitoring (MRM) compatibility. The unique amino acid sequence for the marmoset insulin tryptic peptide was confirmed by running a basic local alignment search tool (BLAST) search and did not share similarities with any common interferences. The intact marmoset insulin peptide differed from human insulin at 5 amino acids and the MITP differed from the same tryptic fragment for human insulin by 1 amino acid (GFFYTPK in human). The MITP was simple to synthesize and, due to its moderate hydrophobicity, made it highly compatible with MRM detection (Supplementary Figure 1). Heavy labeled marmoset insulin tryptic peptide (GFFYAPK (^13^C, ^15^N)) was used as an IS. For each peptide and IS, collision-induced dissociation products of multiple charged precursors were detected by MRM mass spectrometry operating in positive ion mode using ESI. The electrospray voltage, source temperature, and ion source gases 1 and 2 were 5500V, 650°C, 35 units, and 60 units, respectively. For MS/MS optimization, 1 µg/ml solutions of MITP and MITP-IS were made in 20:80 acetonitrile:water and were directly infused at a flow rate of 10µl/min. The most intense isotopic peak of the MH_2_^+2^ ion of the marmoset insulin tryptic peptide and heavy labeled peptide was used as the parent peak. This also corresponded to the data provided by the manufacturer of the fragment (Genscript, Thermo). The most intense product ions that gave a higher m/z than that of the parent ion were chosen for quantifier and qualifier peaks. MRM transitions and charge states for each peptide fragment and IS are summarized in Table 1. Representative chromatograms for the marmoset insulin tryptic peptide, the internal standard, and blank are shown in Figure 1 and representative spectra of the MITP are presented in Supplementary Figure 2.

**Table 1.** Optimized MS/MS parameters for marmoset insulin tryptic peptide (MITP) and the internal standard (IS).

| Compound Name | Parent (m/z) | Daughter (m/z) | Dwell time (ms) | DP (v) | CE (v) | Ion Type |
| --- | --- | --- | --- | --- | --- | --- |
| Quantitative | 415.4 | 625.3 | 195 | 82 | 16 | y<sub>5</sub> Tryptic Peptide |
| Qualifier | 415.4 | 478.3 | 195 | 82 | 17 | y<sub>4</sub> Tryptic Peptide |
| IS | 419.4 | 633.4 | 195 | 82 | 15.65 | y<sub>5</sub> |

**Figure 1.**
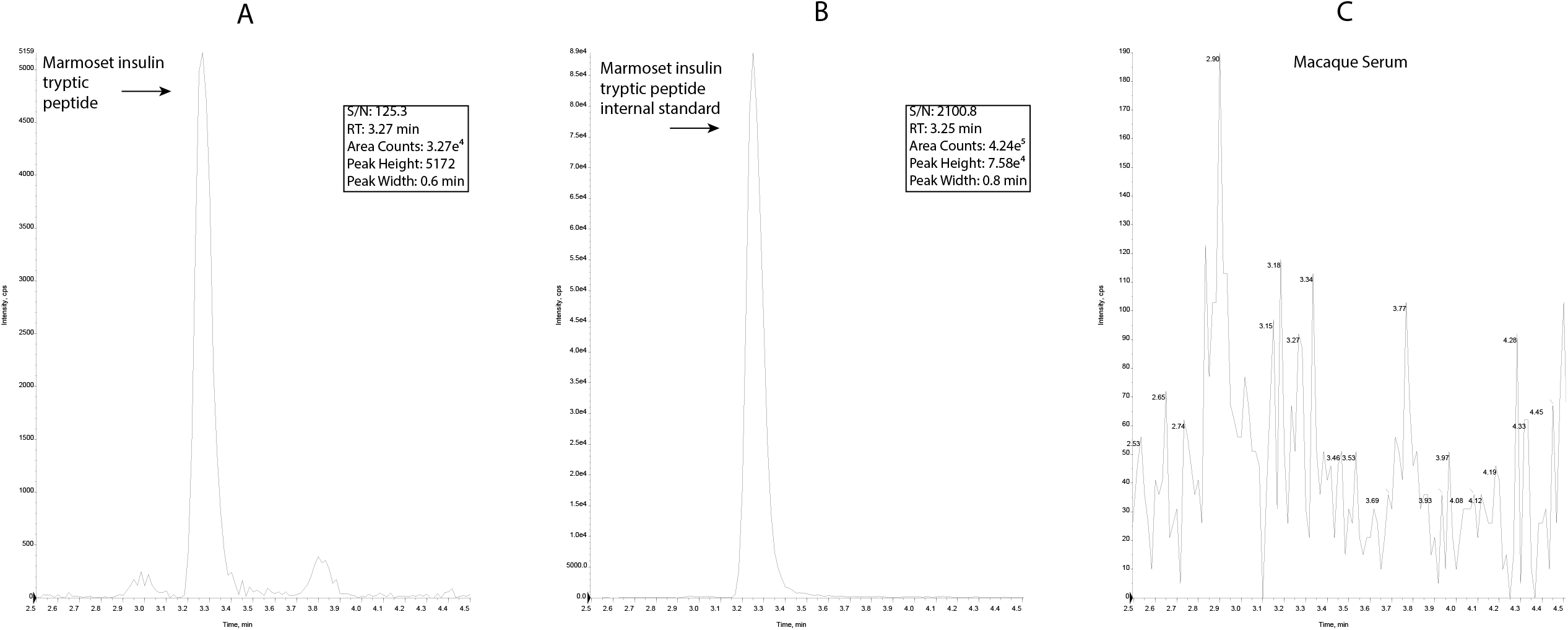
Representative chromatograms for (A) low quality control pool of marmoset insulin tryptic peptide (MITP) in marmoset serum (626 pmol/L), (B) marmoset insulin IS in macaque serum, and (C) macaque serum. The expected retention time for MITP is 3.27 min.

Chromatographic conditions were initially chosen based on the hydropathy of the target peptide; since it was hydrophobic, a low starting concentration of solution B 10 to 30% B was chosen. As expected, the peptide eluted from the column within the linear gradient, however at 50°C, the peptide eluted earlier than intended at 1.7 min and peak splitting was observed. The column oven was reduced to 35°C and the peak eluted later in the gradient (3.27 min) with no splitting.

### Assay Validation

#### Linearity, limit of detection (LOD), and limit of quantitation (LOQ)

The ratio of the peak area of the analyte to the IS was used to calculate the concentrations for the calibration curve. A weighted linear model (1/x) with linear regression was used. The calibration curve had a range of 3100 pmol/L to 24.2 pmol/L over eight points. The calibration range was based on the concentration of human insulin measured by LC-MS/MS [15], but started at a higher concentration based on preliminary data from a marmoset insulin pool. Accuracy of calibration was within 5% at each point measured over 5 runs, including the lowest calibrator. Calibration data is presented in Figure 2. Biological validation data presented below showed that the range of calibration required for the physiological range of marmoset insulin was well above the LOQ. QC pools containing intact marmoset insulin and MITP spiked into macaque serum were used to calculate the LOD and LOQ. The LOD for the method was determined by assaying diluted marmoset serum in 5 replicates to measure an LOD of 4.28 ± 1.05 pmol/L which had a CV of 24.48%. This was confirmed by spiking MITP at a concentration of 6.05 pmol/L and assaying 5 times. The accuracy was 87% ± 29.8%. Similarly, the LOQ was calculated based on measurement of the diluted QC pool 5 times and was 8.62 ± 1.15 pmol/L with a CV of 13.29%. The accuracy of 12.1 pmol/L of MITP spiked into macaque serum was 110% ± 22%.

**Figure 2.**
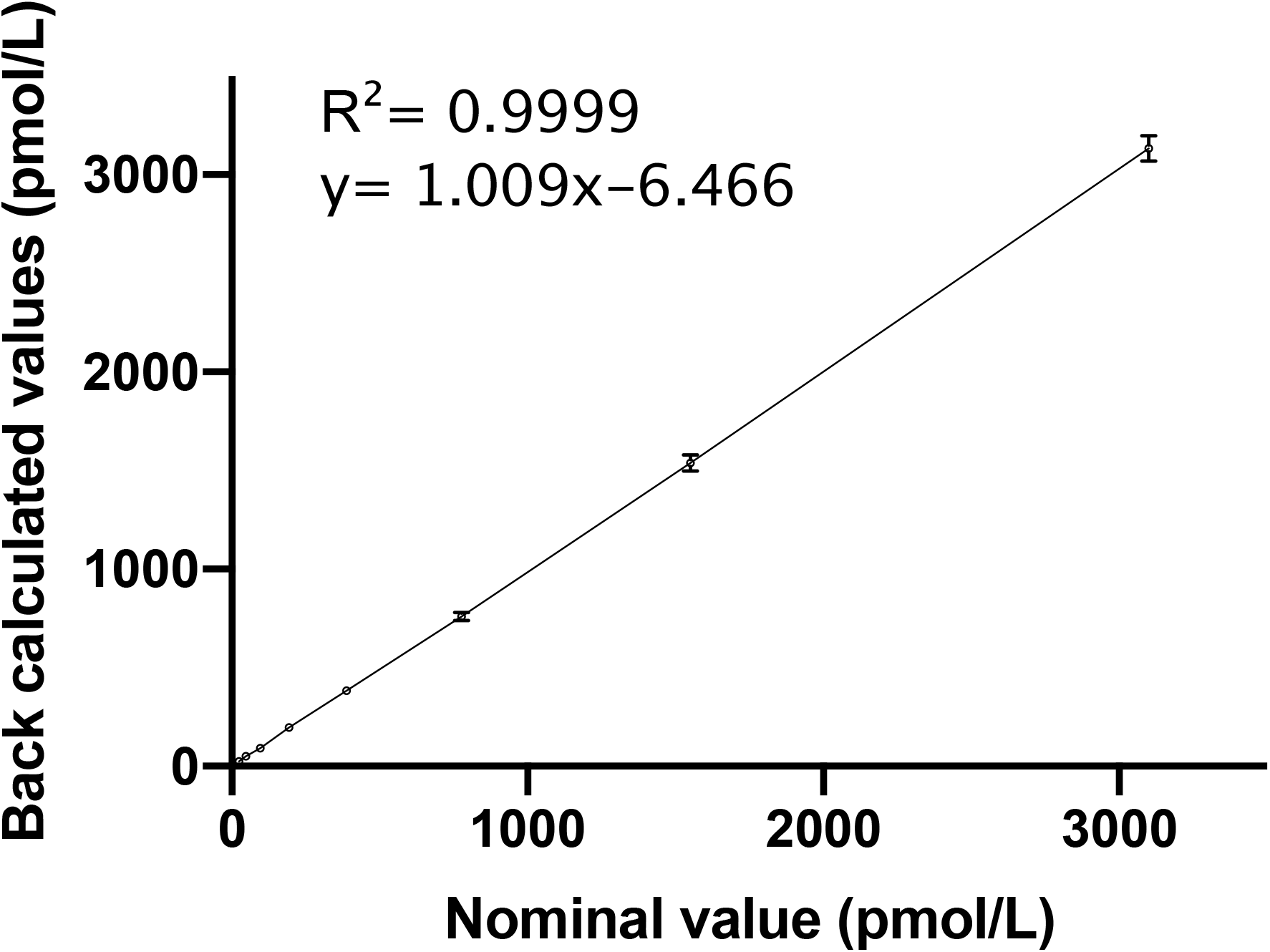
Calibration plot for marmoset insulin tryptic peptide (MITP) in eight determinations from 3100 pmol/L to 24.2 pmol/L. A weighted linear model (1/x) with linear regression was used to calculate the calibration curve. Line equation was y=1.009x-6.466, r^2^ = 0.9999.

#### Precision

Precision was evaluated by measuring the CV of the measurements of the high and low QC pool (Table 2). The intra- and inter-assay CVs for both QC pools were within acceptable range.

**Table 2.** Marmoset insulin quality control (QC) pool data.

| QC pool | Calculated concentration of marmoset insulin (pmol/L ± SD) | Intra-assay CV (# batches) | Inter-assay CV (n replicates) |
| --- | --- | --- | --- |
| Low | 626 ± 59.8 | 2.04%-10.2% (3) | 9.55% (22) |
| High | 1304.16 ± 92.5 | 2.66%-5.78% (3) | 7.09% (23) |

#### Dilutional linearity

A reference preparation of intact marmoset insulin was not available. In order to definitively prove that the method was measuring marmoset insulin, it was purified directly from marmoset pancreas and used in dilutional linearity experiments in macaque serum. The lyophilized material was diluted 0-10,000 times. The back-calculated values were determined based on the calibration curve and the r2 value was 0.9999 (Table 3). The 0 (neat) and 1:100 dilutions were beyond the range of the calibration curve and the concentrations were extrapolated.

**Table 3.** Purified intact marmoset insulin spiked into macaque serum and diluted to generate a linear regression (r^2^ = 0.9999).

| Dilution | Peak area counts | Back calculated concentration (pmol/L ± SD) |
| --- | --- | --- |
| Neat | 2.34E+07 | 539,333 ± 26,102 |
| 1:100 | 2.76E+05 | 5840 ± 537 |
| 1:1000 | 2.14E+04 | 519.3 ± 35.85 |
| 1:10000 | 2.45E+03 | 56.26 ± 5.91 |

### Biological validation

#### Oral glucose tolerance testing

The basal levels of glucose and insulin were 111-144 mg/dL and 128-628pmol/L, respectively. One-way (time) repeated measures ANOVA was significant (p=0.0003) and demonstrated that glucose levels rose significantly above baseline by 15 mins and stayed high until 60 mins. By 120 mins, the glucose levels were indistinguishable from baseline. Overall, insulin concentrations demonstrated an effect of time (p= 0.0037) and post hoc analysis demonstrated there was a trend towards insulin being higher than baseline at 15 mins (p=0.0832) and 30 mins (p=0.06). The percent change in glucose and insulin from baseline are shown in Figure 3. This data also provided the physiological range of marmoset insulin since this was the first time that marmoset insulin was measured directly. This range was 103.8 to 1222 pmol/L.

**Figure 3.**
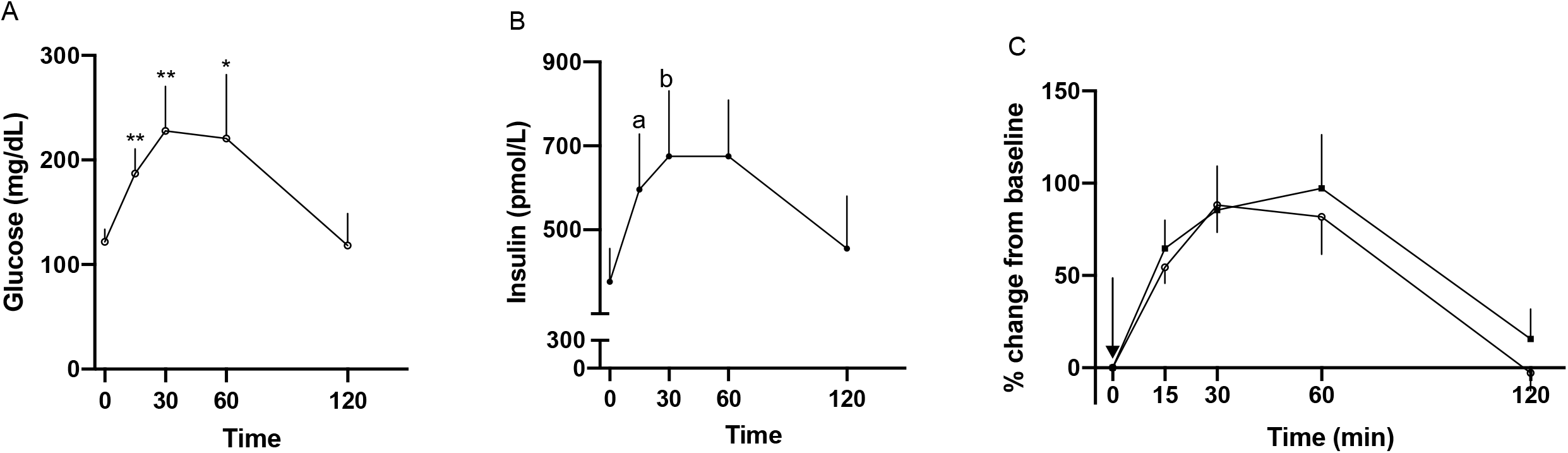
Marmoset serum response to oral glucose tolerance testing (OGTT; n=6). A) glucose (open circles), B) insulin (closed squares), and C) glucose and insulin percent change from baseline. Arrow indicates oral administration of dextrose. * indicates p < 0.05, ** indicates p < 0.01, a indicates p=0.0832, b indicates p=0.06.

#### Comparison with commercial RIA

There was a strong correlation between the previously validated porcine RIA for marmoset insulin and the current method (Figure 4, r = 0.82, p < 0.0001). Based on the LC-MS/MS data, the physiological range was 84.5 to 1164 pmol/L, similar to the results of the OGTT experiment.

**Figure 4.**
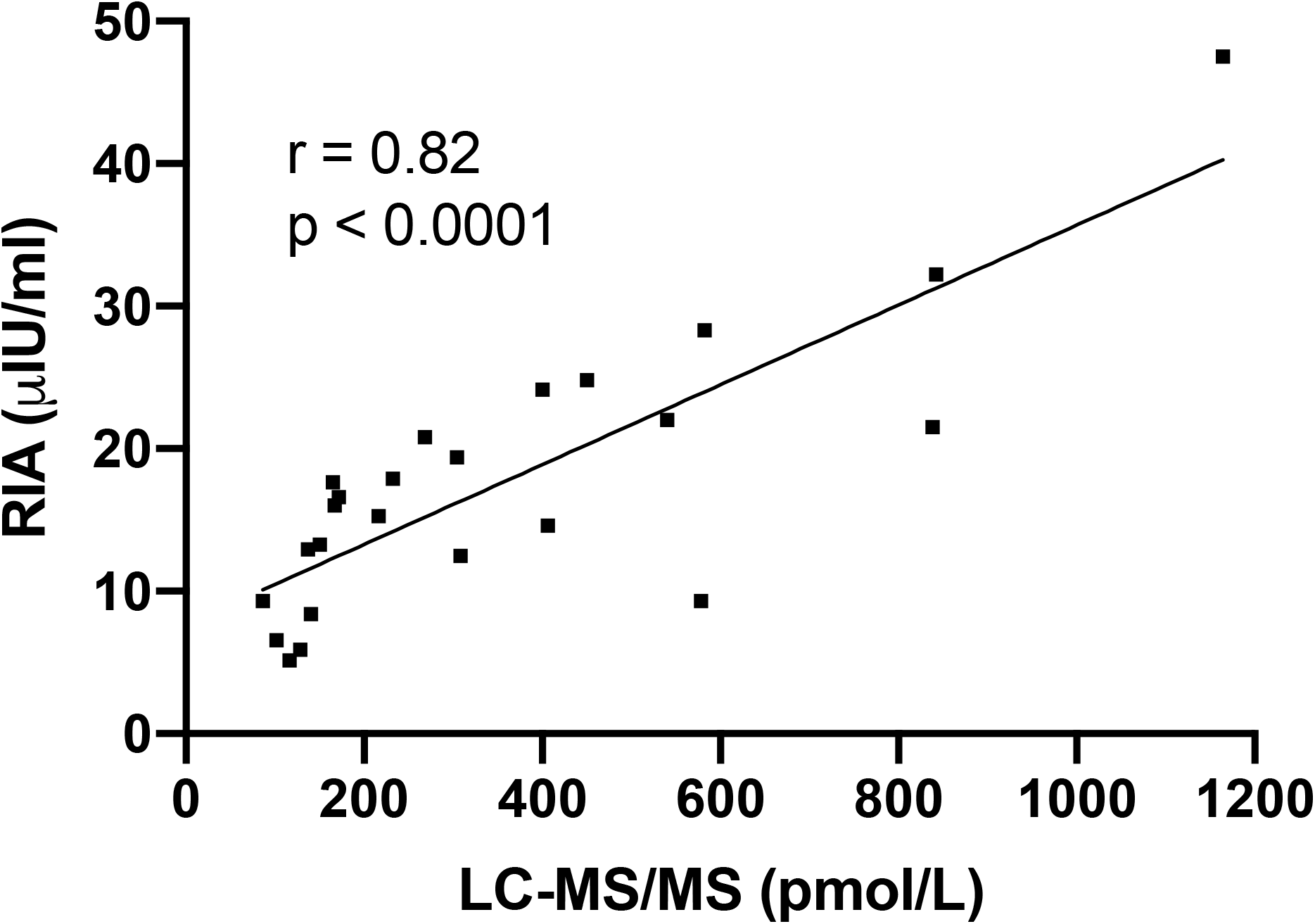
Comparison of 24 marmoset serum samples that were run by a human insulin radioimmunoassay in 2016 and rerun with the newly developed LC-MS/MS method. The correlation between the methods was high (r=0.82, p<.0001).

## DISCUSSION

This paper presents a novel LC-MS/MS based method utilizing a synthetic species-specific tryptic peptide to measure insulin in marmoset serum. The validations demonstrated that the method is reproducible, precise, and accurate according to the guidelines of FDA [21]. Biological validation with OGTT confirmed the expected changes in insulin and glucose levels reported earlier [16] and the concentration changes of glucose and insulin were comparable to humans [23]. The OGTT provided evidence that the method can reliably measure changes in insulin level during physio-pathological conditions. For the second part of the biological validation, high correlation between values obtained for samples previously run using the validated porcine RIA [16] and the current method provided further reliability of the method. To our knowledge, this is now currently the only method to measure marmoset insulin available.

We opted to measure a tryptic peptide of marmoset insulin rather than the intact B chain of the peptide as was done in human studies [15], [19] for two reasons. First, the B chain was harder to synthesize and more expensive than a tryptic peptide. It would have also been difficult to purchase a heavy labeled version of the marmoset B chain for IS. Human studies circumvent this issue by using bovine B chain as the IS, but due to the uniqueness of the structure of the marmoset B chain and that external calibrators and QC material was not readily available, we deemed it critical to have a heavy labeled version of the exact peptide that was being measured for this study. Second, in order to fully develop a marmoset model of aging and disease, there are other important biomarkers that require assay development. The use of trypsin to digest the peptides creates predictable tryptic fragments [17] making this technology adaptable for other metabolites. Furthermore, the simple sample preparation using the µ-elution mixed-mode cation exchange solid-phase extraction plate is amenable to inclusion of additional analytes, which is important due to the low blood volume of marmosets.

There are a number of studies that have been conducted that have measured marmoset insulin and they have exclusively used the discontinued RIA from Millipore that was validated for marmosets [16]. In the Ziegler study, OGTT and stimulation of marmoset pancreas *in vitro* demonstrated the biological validity of the assay. Parallelism and accuracy were also demonstrated. The range of the insulin response for the OGTT and the pancreatic stimulation experiments was approximately 2.5 to 80 µIU/mL. In a study conducted to understand the impact of early life obesity on metabolic parameters, at 12 months of age, obese animals (16.45 + 3.04 µIU/mL) had significantly higher insulin levels than the control animals (1.01 + 0.25 µIU/ mL) [7]. Another study conducted to look at the fasting insulin concentration in female marmosets to understand the combined and independent effects of aging and obesity on metabolism in marmosets showed that the range of insulin was from approximately 2 to 110 3.04 µIU/mL [8]. Before the porcine RIA was validated for marmoset insulin, Wachtman et al., conducted a study to measure the effect of diet on the development of obesity and T2DM in marmosets. They showed that both a high sugar or high fat diet resulted in profound pancreatic islet hyperplasia, which suggested a compensation for increased insulin requirements, though insulin was not measured [24].

Conventional insulin concentration units (µIU/mL) are based on bioefficiency which is defined as the amount of insulin required to achieve a standard glycemic effect. This number has changed several times based on differences in standards and purity of insulin preparations [25]. The factor to convert to SI units is based on the molecular weight of insulin, which differs among different species. For example, human insulin is 5808 Da, marmoset insulin is 5764 Da, and porcine insulin is 5777 Da [26]. The porcine RIA kit insert [16] did not provide any information about the conversion factor that could be used to convert to SI units so we chose the factor 6 (1 µIU/mL = 6 pmol/L) [25]. With this conversion factor, the range of marmoset insulin measured by RIA (approximately 6 to 600 pmol/L) is lower than the range that we present in this paper (84.5 to 1222 pmol/L).

While the RIA was validated, the cross-reactivity of the porcine antibody with marmoset insulin was not investigated. Comparison of the range of marmoset insulin observed in this study with human insulin are similar. In two studies, human insulin measured by LC-MS/MS was shown to be from 18-1080 pmol/L [15] and 86.08 - 774.8 pmol/L [27]. Based on these data, the RIA could have been underestimating marmoset insulin in serum, especially at high concentrations, but it is not possible to know with certainty as marmoset insulin standard material is not available.

There are two limitations to the method validation. First, it is not possible to do a recovery study in the traditional manner since we did not have precise amounts of intact marmoset insulin. We partially overcame this by spiking relative amounts of purified marmoset insulin at different dilutions and demonstrating the linearity and back-calculated values. This obstacle is part of the second limitation which is that we cannot be certain of the absolute concentrations of marmoset insulin since there is no verified standard material to trace back to. Nonetheless, based on the validation parameters presented, we are confident that we have a specific, reproducible method for measurement of marmoset insulin.

In summary, we have developed the only quantitative LC-MS/MS based assay specific for marmoset insulin. It is not reliant on commercially available antibodies or immunoassays and is easily transferable to other labs that have the capacity to measure peptides by LC-MS/MS.

## ACKNOWLEDGMENTS

Special thanks to the staff at Assay Services and the Scientific Protocol Implementation at the WNPRC and the Mass Spectrometry Facility at the UW-Madison Biotechnology Center.

## SUPPLEMENTARY INFORMATION

**Supplementary Figure 1.**
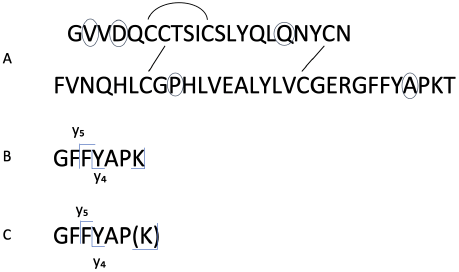
(A) Amino acid sequence for intact marmoset insulin. Disulfide bonds are indicated with the lines connecting the cysteine residues. Circles indicate amino acids that differ from human/macaque insulin. (B) Marmoset insulin tryptic peptide (MITP) and (C) marmoset insulin tryptic peptide IS (MITP-IS) sequences and fragment ions monitored by MRM. Heavy labeled lysine (K) = (^13^C,^15^N).

**Supplementary Figure 2.**
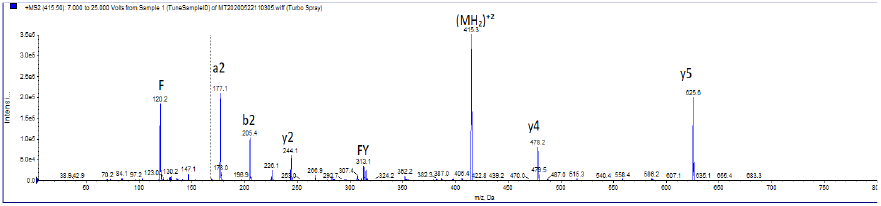
MS/MS spectra of marmoset insulin tryptic peptide (GFFYAPK) MH_2_^+2^ (m/z = 415.3). Collision energy ramped from 7 to 25 volts.

## Notes

### Competing Interest Statement

The authors have declared no competing interest.

